# HTSlib-js: Developer toolkit for a client-side JavaScript interface for HTSlib

**DOI:** 10.1101/397935

**Authors:** Jeongbin Park, Liam H. Childs, Matthias Schlesner, Roland Eils

## Abstract

**Background:** An increasing number of bioinformatics tools are developed in JavaScript to provide an interactive visual interface to users. However, there are no tools yet that enable fast client-side analysis of large, high-throughput sequencing file formats, such as BAM and VCF files.

**Results:** We present HTSlib-js, an asm.js-based JavaScript wrapper for HTSlib, the *de facto* standard for processing BAM and VCF files. HTSlib-js exploits recent technological advances in web browser engines that dramatically increase the performance of browser-based tools to enable swift processing of files in the aforementioned formats. HTSlib-js enables quick development of JavaScript-based applications that include processing of aligned sequence reads and variant calling data.

**Conclusions:** HTSlib-js constitutes a toolkit for developers to easily write fast, accessible, browser-based applications that place an emphasis on data visualization, interactivity and privacy. Real-world examples demonstrate the capabilities of HTSlib-js and serve as guide for own developments.

## Background

The high-throughput sequencing library (HTSlib) and the corresponding data formats [1], were introduced to process data for the 1000 Genome Project [2] and has now become the de facto standard for high-throughput sequencing (HTS) file formats. Many short-read alignment [3–5] and variant calling tools [1, 6, 7] use HTSlib to process sequence reads (SAM, BAM) and variant call data (VCF, BCF).

Tools that depend on HTSlib are typically run from the command-line restricting their accessibility to users who have a specific set of computational skills. To improve accessibility, several tools attempt to provide a better user experience in the form of a web-interface and directly process HTS files in the web-browser [8, 9]. This necessitates the use of JavaScript, the only language supported by all major browsers. However, JavaScript was not initially developed to cater for heavy computational demands and the sophisticated analysis of large files suffers from poor performance in the browser. Tools which perform data processing on the server-side can overcome these limitations, but transfer of large data files can be slow, and transfer of sensitive data like genomic data to central servers might be restricted due to privacy concerns. To overcome JavaScript’s performance issues, the Mozilla foundation recently announced a limited subset of JavaScript called ‘asm.js’ [10]. This subset removed performance intensive features from the language and the resulting code could be highly optimized by the web browser’s JavaScript engines. Moreover, because asm.js shares many similarities with assembly language, it could also be the target of a compiler. Based on this idea, a compiler called ‘Emscripten’ [11] adopted asm.js, allowing it to compile C/C++ code into asm.js. Thanks to further optimization technologies recently introduced to web browsers, the execution speed of asm.js programs written in C/C++ code and compiled using Emscripten is much closer to that of a native binary executable.

In this work, we present HTSlib-js, a software development toolkit providing a port of the original HTSlib that can be easily compiled into asm.js using Emscripten. Along with the toolkit, we also provide real-world examples for developers to easily understand how to access/process supported file formats using HTSlib-js.

## Implementation

### Compile HTSlib-js using Emscripten

To easily configure and build the project, we provide a CMake configuration file. Running CMake via the Emscripten SDK in the source code tree will automatically invoke Emscripten to compile all required files and included examples (Fig. 1A). The CMake script can also be used to generate native binaries (Fig. 1B) because the analysis source code and HTSlib are written in C/C++. This enables developers to provide a native binary of their tools for users who prefer the command line environment or the integration into existing processing pipelines.

**Figure 1.**
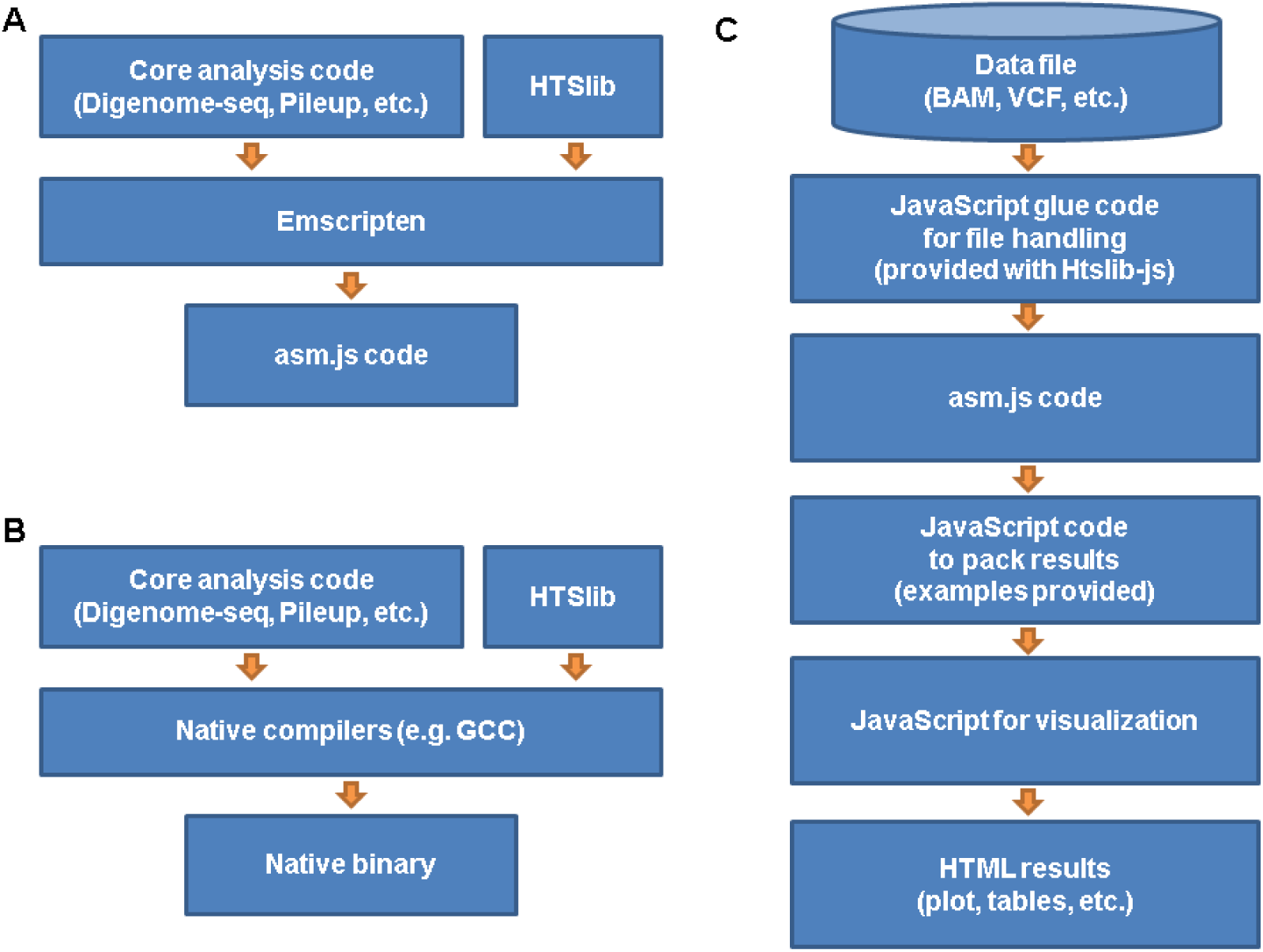
HTSlib-js Workflow. (A) Compilation of analysis tools using HTSlib with Emscripten. (B) Compilation of analysis tools using HTSlib with native compilers. Almost no modification is needed in the analysis code. (C) A possible workflow of a webtool using Emscripten-compiled asm.js code.

### CIGAR string byte alignment

Small aspects of HTSlib needed to be adjusted to enable full compatibility with the Emscripten compiler. The API to retrieve CIGAR information from BAM files (bam_get_cigar) uses unaligned memory access, which is currently not supported by Emscripten. We demonstrate the correct retrieval of the CIGAR string with an example called Digenome-seq.

### Speed optimization

The current Emscripten implementation already supports a native way to read files. However, the native implementation is too slow to process large input files like BAM files. Therefore HTSlib-js implements file reading routine featuring a double buffering strategy to read BAM files more efficiently.

### Examples

In addition to the core library and interface code, the HTSlib-js source tree also contains three real-world examples to work with file formats supported by HTSlib (Fig. 1C); a basic pileup tool, a somatic variant quality control tool and a tool for identifying and visualising double stranded breaks called Digenome-seq [12]. The pileup tool can process BAM/CRAM files to produce a basic pileup similar to that of the samtools command line tool. The quality control tool can process VCF files and produces plots to assess the quality of somatic variants. The third example is the complete working browser-based Digenome-seq program [13]. As described in the previous section, the examples are automatically built by CMake (via Emscripten for JavaScript targets, or normal CMake for native binaries).

Due to limitations imposed by the browser environment, HTSlib-js only supports reading from, but not writing to the hard-drive. Developers wishing to save data to the hard-drive will first need to store results in memory and then prepare a downloadable file using the HTML5 Blob API.

## Results

Using Emscripten, we produced a fully functional HTSlib wrapper that can be used to create JavaScript-based programs capable of quickly processing SAM, BAM, VCF and BCF files. Examples demonstrate the analysis of SAM and BAM files (pileup example), and VCF and BCF files (quality control example) in browser-based tools including the retrieval of CIGAR strings (Digenome-seq). We compared the performance of asm.js binaries in a web browser to native binaries using a 1.3 GiB BAM file. In our speed benchmark, the compiled asm.js binaries showed a performance relatively close to the native binary. There was only a 5x difference in the pileup example and 2x difference in Digenome seq (Table 1), which will be acceptable for many applications. Hence HTSlib-js overcomes a major drawback of earlier browser-based tools.

**Table 1.** Benchmark results of the compiled native vs asm.js binaries.

|  | Pileup | Digenome-seq |
| --- | --- | --- |
| Native binary | 28 sec. | 18 sec. |
| Web browser (Firefox) | 139 sec. | 34 sec. |

Note: The test was performed using a 1.3GiB BAM file. The system used for testing was equipped with an Intel(R) Core(TM) i5-4590 CPU (3.30GHz) running a Linux system.
Firefox on the same system was used for the web benchmark.

Due to the browser environment HTSlib-js based tools are likely to be strong in the following aspects: 1) Visualization. By nature, the web browser is highly visual and many libraries exist that provide easily understandable and optically attractive visualization of scientific data. 2) Interactivity. Many JavaScript visualization libraries place great emphasis on interactivity allowing users to explore their data to assist the generation of new hypotheses. 3) Accessibility. Browser-based tools require no installation or up-dates removing the need for specialized knowledge when obtaining and maintaining a tool. Furthermore, web-browsers provide a familiar environment that many users will be comfortable with. 4) Privacy. Data provided to HTSlib-js based tools are not required to leave the browser environment permitting sensitive data to be analyzed completely within the user’s computer without transfer to a server.

## Conclusions

We provide a software developer’s toolkit called HTSlib-js that will facilitate the development of HTS analysis tools within the browser environment. We show that the library can be used to develop tools that process HTS files at close-to-native speeds. HTSlib-js paves the way for a new generation of fast, accessible, browser-based tools that place an emphasis on data visualization, interactivity and privacy.

## Availability and requirements

### Project name

HTSlib-js

### Project home page

https://github.com/eilslabs/htslib-js

### Operating system(s)

Platform independent

### Programming language

C/C++, ECMAScript (JavaScript)

### Other requirements

Emscripten

### License

GNU LGPL

### Any restrictions to use by non-academics

license needed

## List of abbreviations

SAM: Sequence alignment map
BAM: Binary alignment map
VCF: Variant call format
BCF: Binary variant call format
HTS: High-throughput sequencing
SDK: Software development kit
CIGAR: Concise idiosyncratic gapped alignment report
HTML: Hypertext markup language
API: Application programming interface
GiB: Gibibyte
GNU LGPL: GNU lesser general public licence

## Declarations

### Authors’ contributions

JP designed this project. JP and LHC implemented the software. JP, LHC, MS, and RE wrote the manuscript. MS and RE supervised the project.

